# A nucleocapsid-tagged fluorescent pangolin coronavirus enables authentic-virus neutralization assays and localizes the antiviral activity of cepharanthine to viral entry

**DOI:** 10.64898/2026.08.16.742902

**Authors:** Yuezhen Ma, Shanshan Lu, Shengdong Luo, Yue Hu, Chenyang Wang, Xiao Zhang, Li Deng, Caiwu Li, Weiwei Chen, Wenxin Zheng, Lihua Song

**Affiliations:** College of Life Science and Technology, Beijing University of Chemical Technology, Beijing 100029, China; Senior Department of Infectious Diseases, The Fifth Medical Center of Chinese PLA General Hospital, National Clinical Research Center for Infectious Diseases, 100073 Beijing, China; College of Veterinary Medicine, Xinjiang Agricultural University, Urumqi 830052, China; The 82nd group army hospital of the PLA, Baoding, China; China Conservation and Research Centre for the Giant Panda, Key Laboratory of State Forestry and Grassland Administration on the Giant Panda, Cheng du 610066, Sichuan Province, China

**Keywords:** pangolin coronavirus, GX_P2V, reporter virus, nucleocapsid protein, mBaoJin, reverse genetics, neutralizing antibody, cepharanthine, virus entry, time-of-addition assay

## Abstract

Replication-competent coronaviruses carrying a fluorescently tagged structural protein are rare, and the only well-developed example tags the membrane (M) protein. Using the highly attenuated pangolin coronavirus GX_P2V(short_3UTR) as a backbone, we assembled a full-length cDNA clone by multifragment in vitro ligation and recovered GX_P2V-mBJ-N, a recombinant virus in which the bright monomeric green fluorescent protein mBaoJin is fused to the N terminus of the nucleocapsid (N) protein. The equivalent C-terminal fusion was not viable. GX_P2V-mBJ-N is infectious in Vero E6 cells but is attenuated and genetically unstable in the absence of exogenous N; supplying wild-type N in trans allows propagation to high titer with the fusion intact. Because mBaoJin is translated as part of N, fluorescence reports viral antigen synthesis directly, and infection can be quantified in a plate reader without immunostaining or plaque reduction. We applied the reporter virus in two settings. First, sera from ten healthy blood donors were titrated in parallel against GX_P2V-mBJ-N and a vesicular stomatitis virus (VSV)-based GX_P2V pseudovirus. The two assays agreed on rank order, but pseudovirus titers were about 10-fold higher overall and diverged by at least 70-fold in one donor, showing that the two formats are not interchangeable for individual samples. Second, time-of-addition experiments identified entry as the principal target of cepharanthine (EC_50_, 0.20 μM under continuous treatment), whereas preincubation of virions with 25 μM drug had no effect, excluding direct virion inactivation. GX_P2V-mBJ-N provides a practical authentic-virus platform for coronavirus serology and antiviral pharmacology.

**IMPORTANCE:** Fluorescent reporter coronaviruses are normally built by placing a free fluorescent protein in an accessory gene or by releasing it from the nucleocapsid gene with a 2A peptide, so the marker is a soluble cytoplasmic protein rather than part of the virion. Fusing a fluorescent protein to a structural protein is much harder, and until now the only well-developed example tagged the M protein of a murine coronavirus. We show that the N terminus of the coronavirus N protein also tolerates such a fusion, in a SARS-CoV-2-related virus, provided that wild-type N is supplied in trans. Because the tag is built into the most abundant viral protein, fluorescence becomes a direct quantitative readout of viral antigen synthesis, which turns technically demanding authentic-virus assays into plate-reader measurements. The backbone is a highly attenuated, non-pathogenic pangolin coronavirus, so these assays are accessible to laboratories without high-level containment.

## INTRODUCTION

Coronaviruses (family *Coronaviridae*) are enveloped, positive-sense, single-stranded RNA viruses whose genomes of 26 to 32 kb are the largest known among RNA viruses (1, 2). Three highly pathogenic human coronaviruses have emerged since 2003: SARS-CoV-1, MERS-CoV, and SARS-CoV-2 (3–5). The continuing diversification of Omicron sublineages, which combine efficient transmission with pronounced immune evasion, has sustained the demand for efficient coronavirus reverse genetics systems and for robust antiviral and serological assays (6–8).

Full-length coronavirus cDNA clones are assembled either by multifragment in vitro ligation or as bacterial artificial chromosomes (9, 10). Reporter derivatives of SARS-CoV-2 have been built on both platforms, for example by inserting a Venus-2A cassette upstream of the nucleocapsid (N) gene so that 2A-mediated ribosomal skipping releases free Venus alongside functional N (11), or by replacing ORF7 with mNeonGreen (12). In all such designs the fluorophore is a soluble cytoplasmic marker and is not a component of the virion. Viruses in which a fluorescent protein is physically fused to a structural protein would additionally report particle localization, assembly, and egress, but they remain scarce, because structural proteins tolerate insertions poorly. The one well-developed example is a murine hepatitis virus carrying a GFP-tagged membrane (M) protein, which assembles only when wild-type M is supplied in trans (13). No comparable virus has been reported for a SARS-CoV-2-related coronavirus, and the N protein, which is the most abundantly expressed viral protein in infected cells and the one whose synthesis most closely tracks the progress of infection, has not been successfully tagged in a replicating coronavirus.

We previously described a SARS-CoV-2 surrogate model based on the pangolin coronavirus GX_P2V(short_3UTR), a cell culture-adapted, highly attenuated live-vaccine candidate that carries a spontaneous 104-nucleotide deletion in the 3′ untranslated region (3′ UTR) (14–16). This strain causes no disease in immunocompetent animals yet elicits cross-protective immunity against SARS-CoV-2. Using a vesicular stomatitis virus (VSV)-based GX_P2V pseudovirus, we found that Chinese vaccinees who experienced Omicron breakthrough infection after control measures were lifted in early 2023 carried high-titer cross-neutralizing antibodies against GX_P2V (17). Separately, drug-repurposing screens in the GX_P2V(short_3UTR) infection model identified cepharanthine (CEP) as a potent inhibitor, although the step of the replication cycle at which it acts was not defined (18).

Here we used the GX_P2V(short_3UTR) genome as a backbone to construct GX_P2V-mBJ-N, a replication-competent recombinant virus in which the green fluorescent protein mBaoJin is fused directly to the N protein. We characterized its replication kinetics, genetic stability, and dependence on N complementation, and then applied it to two problems for which an authentic-virus readout matters: a head-to-head comparison of neutralizing titers measured with the reporter virus and with the VSV pseudovirus in sera from healthy blood donors, and a time-of-addition analysis that localizes the antiviral activity of CEP. Because mBaoJin is translated as part of N, fluorescence intensity is a direct, plate-reader-compatible measure of viral antigen synthesis, which simplifies work with authentic virus.

## RESULTS

### Design and assembly of a full-length GX_P2V(short_3UTR) cDNA encoding an mBaoJin-N fusion

The genome of GX_P2V(short_3UTR) is 29,729 nucleotides long (Fig. 1A). We assembled it from five cDNA fragments joined by in vitro ligation. Fragment junctions were placed at natural or engineered type IIS restriction sites, so that digestion generates unique cohesive ends without altering the viral sequence. A T7 promoter and a poly(A)_30_ tract were appended to the 5′ and 3′ ends, respectively, so that the in vitro transcript would resemble authentic genomic RNA (Fig. 1B).

**FIG 1.**
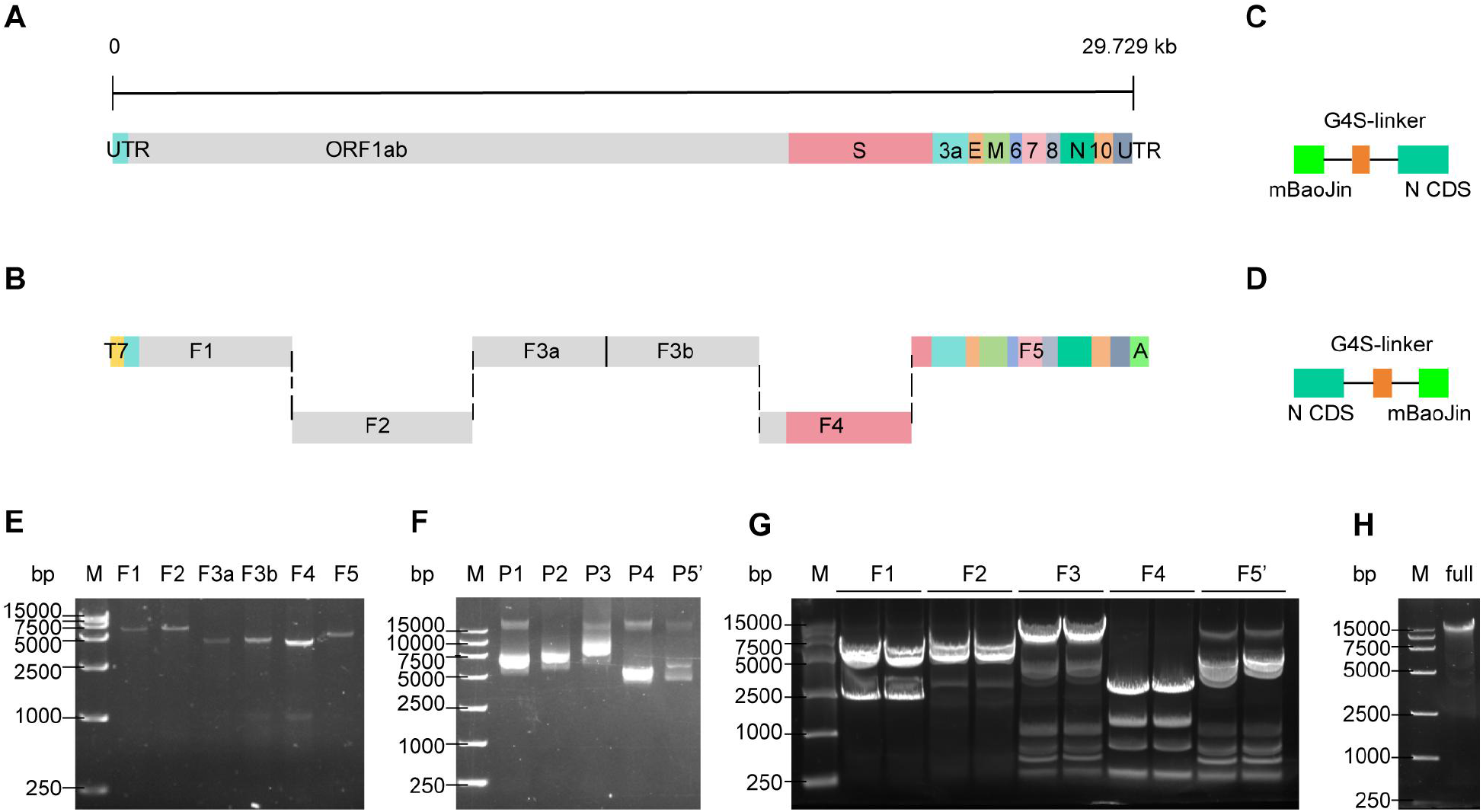
Design and in vitro assembly of the GX_P2V-mBJ-N reverse genetics system. (A) Genome organization of GX_P2V(short_3UTR), showing the open reading frames and the 5′ and 3′ untranslated regions (UTRs). (B) Assembly strategy. A T7 promoter (yellow) and a poly(A) tract (green) were added at the 5′ and 3′ ends of the genome, respectively, and the full-length cDNA was divided into five fragments (F1 to F5) for in vitro ligation. (C and D) The two fusion designs, with mBaoJin joined to the N terminus (C) or the C terminus (D) of the N protein through a flexible G4S linker. (E) The five PCR fragments resolved on a 0.8% agarose gel: F1, 6,237 bp; F2, 6,216 bp; F3a, 4,289 bp; F3b, 4,291 bp; F4, 3,958 bp; and F5, 4,874 bp. Marker range, 250 to 15,000 bp. (F) The five recombinant plasmids resolved on a 0.8% agarose gel. F3a and F3b were cloned into a single vector, and plasmid 5 was modified to give plasmid 5′, which carries the mBaoJin insertion. (G) Restriction digests with BsaI, Esp3I, and BciVI resolved on a 0.8% agarose gel; all bands matched the predicted sizes. (H) The assembled full-length GX_P2V-mBJ-N cDNA.

To label the virion itself, we fused mBaoJin to either the N or the C terminus of the N protein through a flexible G4S linker (Fig. 1C and D). Fusion to the N protein has a second advantage: it couples fluorescence output to viral antigen synthesis rather than to an independently translated marker. We chose mBaoJin, a monomeric derivative of StayGold, because it is 70% to 140% brighter than eGFP and is unusually photostable and chemically stable (19). Viral RNA was reverse transcribed, and the five fragments were amplified with a high-fidelity polymerase (Fig. 1E), cloned individually into pET-40b(+) (Fig. 1F), and released by digestion with BsaI, Esp3I, and BciVI (Fig. 1G). Ligation with T4 DNA ligase yielded full-length cDNA of the expected size (Fig. 1H).

### Recovery of GX_P2V-mBJ-N and its propagation by N complementation

The coronavirus N protein increases the infectivity of genomic RNA, and the two fusion configurations (mBJ-N and N-mBJ) were expected to differ in how well they preserved N function. Because the N proteins of pangolin-CoV and SARS-CoV-2 share 97.8% amino acid identity, we expected the heterologous SARS-CoV-2 N to complement the fusion constructs (20). We therefore electroporated the two genome-length transcripts separately into Vero cells stably expressing wild-type SARS-CoV-2 N (N-Vero cells) and harvested the supernatant at 48 h as passage 0 (P0). Fresh N-Vero cells inoculated with P0 of the mBJ-N construct became green fluorescent on day 3, and the signal intensified by day 4; this supernatant was designated P1. The N-mBJ construct never produced fluorescence, indicating that the C terminus of N tolerates insertion poorly.

Fluorescence increased only modestly over eight passages in N-Vero cells (Fig. 2A), which we attribute to the low level of N expressed by that line. Passage in hACE2-293T cells transiently transfected with an N expression plasmid gave a much stronger signal by P11 (Fig. 2B), showing that the tagged virus can be produced at scale when N is supplied in trans. P11 virus was aliquoted, titrated, and stored for all subsequent experiments.

**FIG 2.**
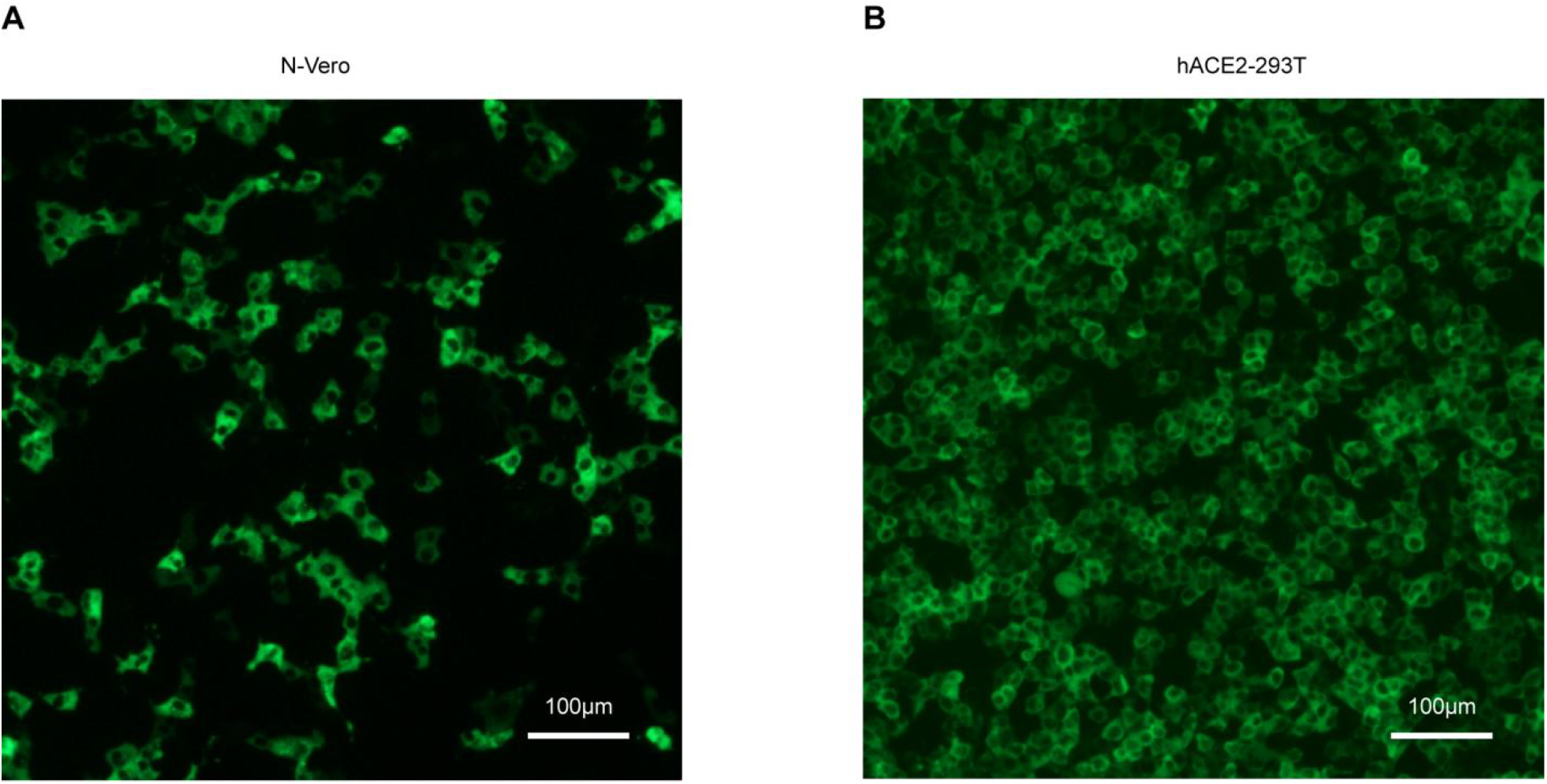
Recovery and passage of the fluorescent reporter virus GX_P2V-mBJ-N. Fluorescence of GX_P2V-mBJ-N in N-Vero cells at passage 8 (A) and in hACE2-293T cells transiently expressing N at passage 11 (B).

The rescued virus, designated GX_P2V-mBJ-N, also infected parental Vero E6 cells that do not express exogenous N (Fig. 3). RT-PCR across the junction followed by Sanger sequencing confirmed an intact mBaoJin-N fusion gene.

**FIG 3.**
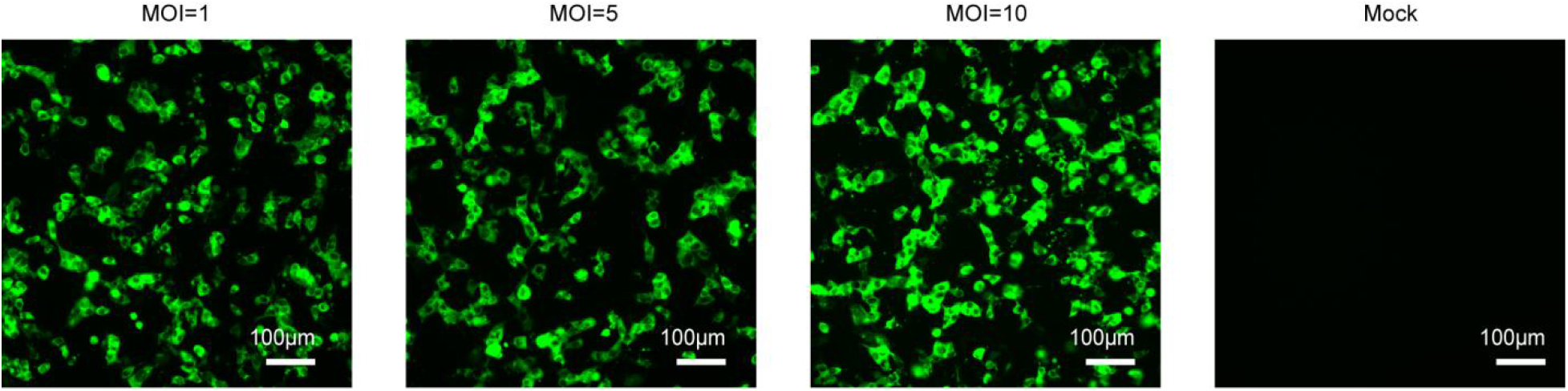
Infection of Vero E6 cells by GX_P2V-mBJ-N. Fluorescence micrographs of Vero E6 cells infected at MOIs of 1, 5, and 10.

### GX_P2V-mBJ-N is infectious but attenuated and genetically unstable without N complementation

A fluorescent protein appended to N could interfere with nucleocapsid assembly, so we compared GX_P2V-mBJ-N with the parental GX_P2V(short_3UTR) (hereafter GX_P2V-WT) in parental Vero E6 cells for replication kinetics, genetic stability, and plaque formation. Multistep and single-step growth curves were obtained at MOIs of 0.01 and 5, respectively (Fig. 4). At an MOI of 0.01 the two viruses were indistinguishable early in infection, but after 36 h the reporter virus fell behind: GX_P2V-WT reached approximately 6.8 and 6.9 log_10_ copies/μL at 48 and 72 h, whereas GX_P2V-mBJ-N reached only about 5.7 log_10_ copies/μL at 72 h (Fig. 4A). At an MOI of 5, GX_P2V-mBJ-N again replicated more slowly through the exponential phase and accumulated fewer genomes, although the difference at 72 h was smaller (7.4 versus 6.5 log_10_ copies/μL) (Fig. 4B).

**FIG 4.**
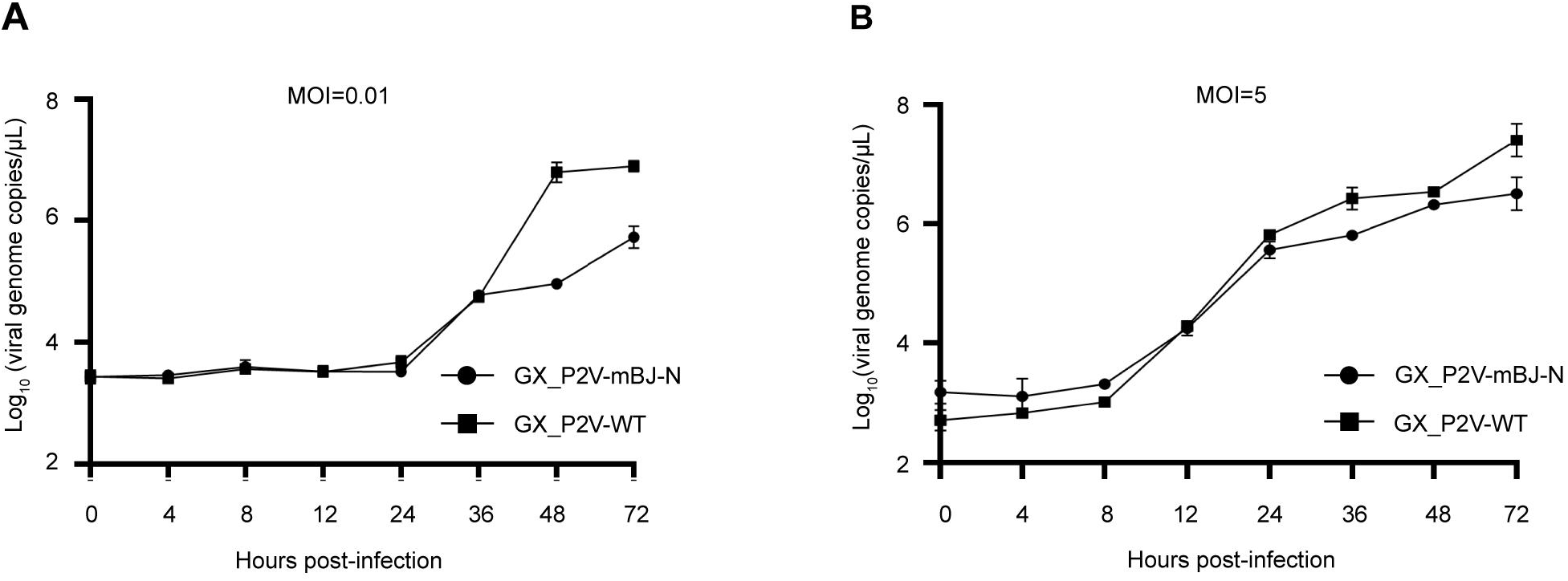
Replication kinetics of GX_P2V-mBJ-N and GX_P2V-WT. Vero E6 cells were infected at an MOI of 0.01 (A) or 5 (B). Viral genome copy numbers in culture supernatants were determined by RT-qPCR at the indicated times postinfection using a plasmid standard curve (R^2^ = 0.9956; plasmid standard, 1.17 × 10^0^ to 1.17 × 10^7^ copies per reaction). Data are means ± standard deviations from three independent experiments.

Genetic stability was assessed over four passages in parental Vero E6 cells (Fig. 5). Green fluorescence was brightest and most widespread at Vero-P1, fell sharply at Vero-P2, and was barely detectable at Vero-P4, while cell morphology in bright-field images remained normal (Fig. 5A). The 902-bp product spanning the mBaoJin-N junction weakened progressively from Vero-P1 to Vero-P4 (Fig. 5B), and by Vero-P4 an internal deletion had reduced the insert to a 141-bp mBaoJin remnant at the N terminus of N (Fig. 5B and C). Western blotting of infected Vero cells detected the full-length fusion protein (predicted mass, 72.2 kDa) together with a species of approximately 51 kDa at Vero-P1, a size intermediate between wild-type N (45.6 kDa) and the fusion protein, but only the 51-kDa species at Vero-P4 (Fig. 5D). This species matches the product of the 141-bp remnant retained at Vero-P4, with 47 residual amino acids (∼5 kDa) appended to wild-type N. The smaller species was thus present as early as Vero-P1, indicating that the deletion arises during early passage in N-Vero cells and that the truncated variant rapidly outgrows the intact reporter virus once N is no longer supplied in trans.

**FIG 5.**
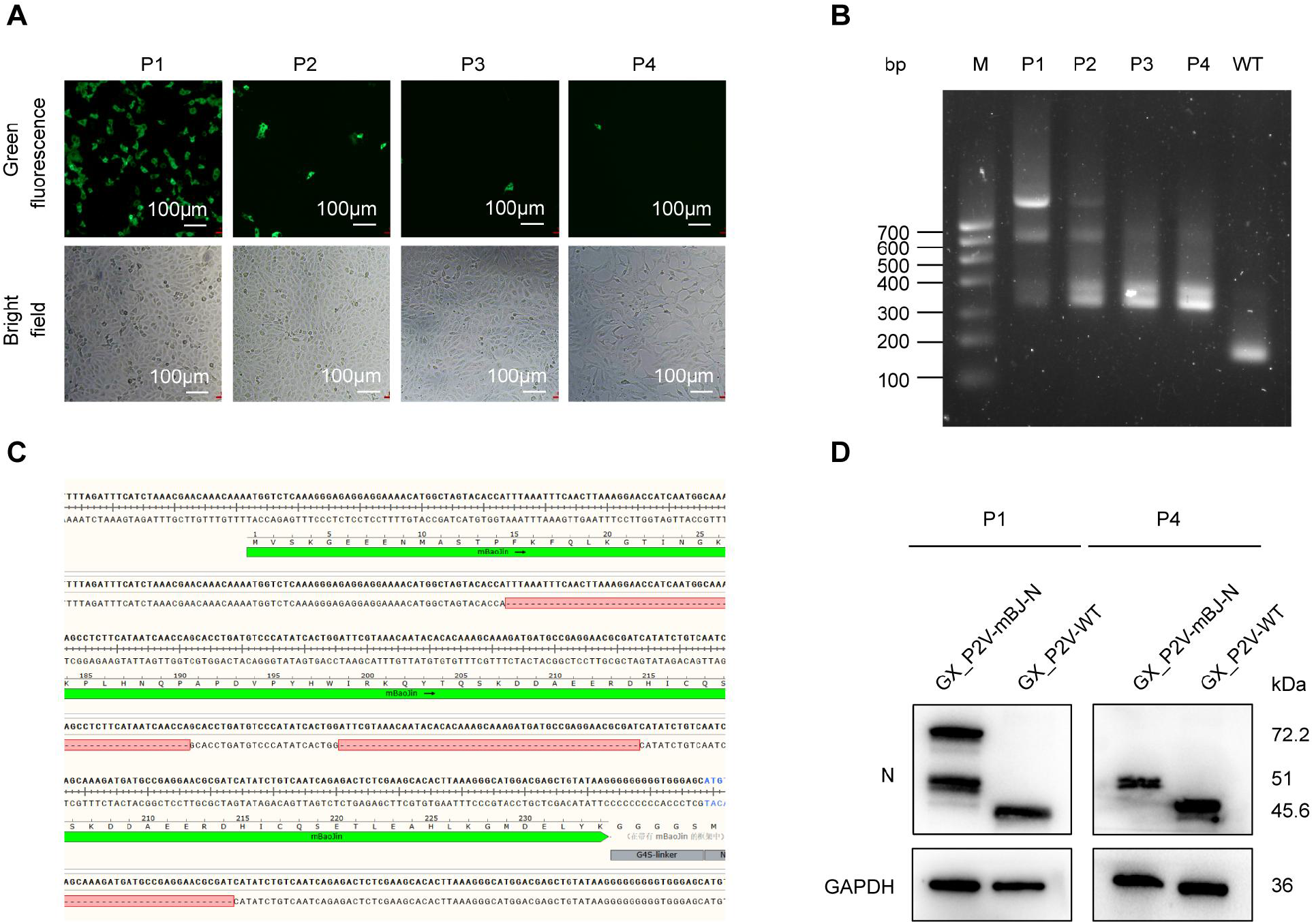
Serial passage of GX_P2V-mBJ-N in Vero E6 cells (P1 to P4). (A) Green fluorescence (top) and bright-field (bottom) micrographs of GX_P2V-mBJ-N at passages 1 to 4. (B) RT-PCR products from GX_P2V-WT and GX_P2V-mBJ-N (P1 to P4). The amplicon spans the full-length mBaoJin gene (702 bp) plus 100 bp of flanking sequence on each side; the WT template yields only the 200-bp flanking product. Marker range, 100 to 700 bp. (C) Sequence alignment of GX_P2V-mBJ-N at passages 1 and 4; deleted sequences are boxed in red. (D) Western blot of the mBaoJin-N fusion protein at passages 1 and 4. Vero E6 cells were infected with GX_P2V-mBJ-N (P1 or P4) or GX_P2V-WT at an MOI of 0.01, and lysates were probed with an antibody against the SARS-CoV-2 N protein. GAPDH (36 kDa) served as the loading control.

Plaque formation was examined in BGM cells. Unlike GX_P2V-WT, neither Vero-P1 nor Vero-P4 reporter virus produced detectable plaques (Fig. 6). Because both preparations infect cells and accumulate viral genomes, the loss of plaques indicates that tagging N impairs the efficiency of viral spread rather than abolishing replication, which is consistent with the slower growth kinetics shown in Fig. 4 and with the requirement for N in trans.

**FIG 6.**
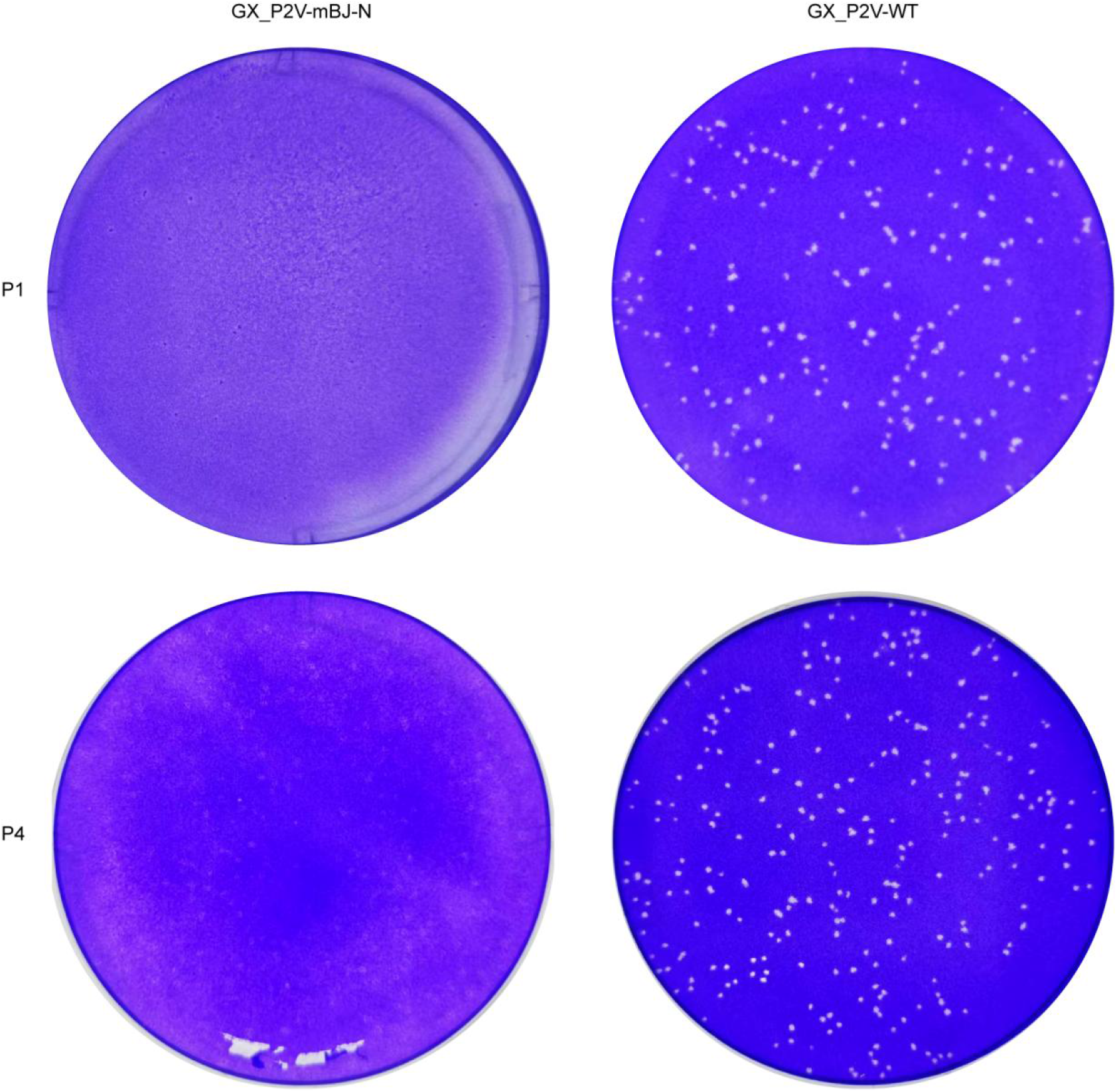
Plaque phenotype of GX_P2V-WT and of GX_P2V-mBJ-N passaged in Vero E6 cells. BGM cells were infected with GX_P2V-mBJ-N (P1 or P4) or GX_P2V-WT, fixed 5 days postinfection, and stained with crystal violet. Representative wells are shown.

### Neutralizing titers measured with the reporter virus and with pseudovirus diverge in individual sera

Because mBaoJin is expressed as part of N, replication of the reporter virus can be quantified as fluorescence intensity in a plate reader, without immunostaining or plaque reduction. Technical difficulty is the main reason that most coronavirus neutralization data are still generated with pseudoviruses; the reporter virus removes that barrier, making an authentic-virus neutralization assay as simple to run as a pseudovirus assay. Using a VSV pseudovirus, we previously reported a geometric mean titer (GMT) of 362 against GX_P2V in vaccinees with Omicron BF.7 breakthrough infection in early 2023 (17). To ask how well pseudovirus titers predict neutralization of the authentic virus, we titrated sera from ten healthy blood donors against the VSV pseudovirus and GX_P2V-mBJ-N in parallel (Fig. 7).

**FIG 7.**
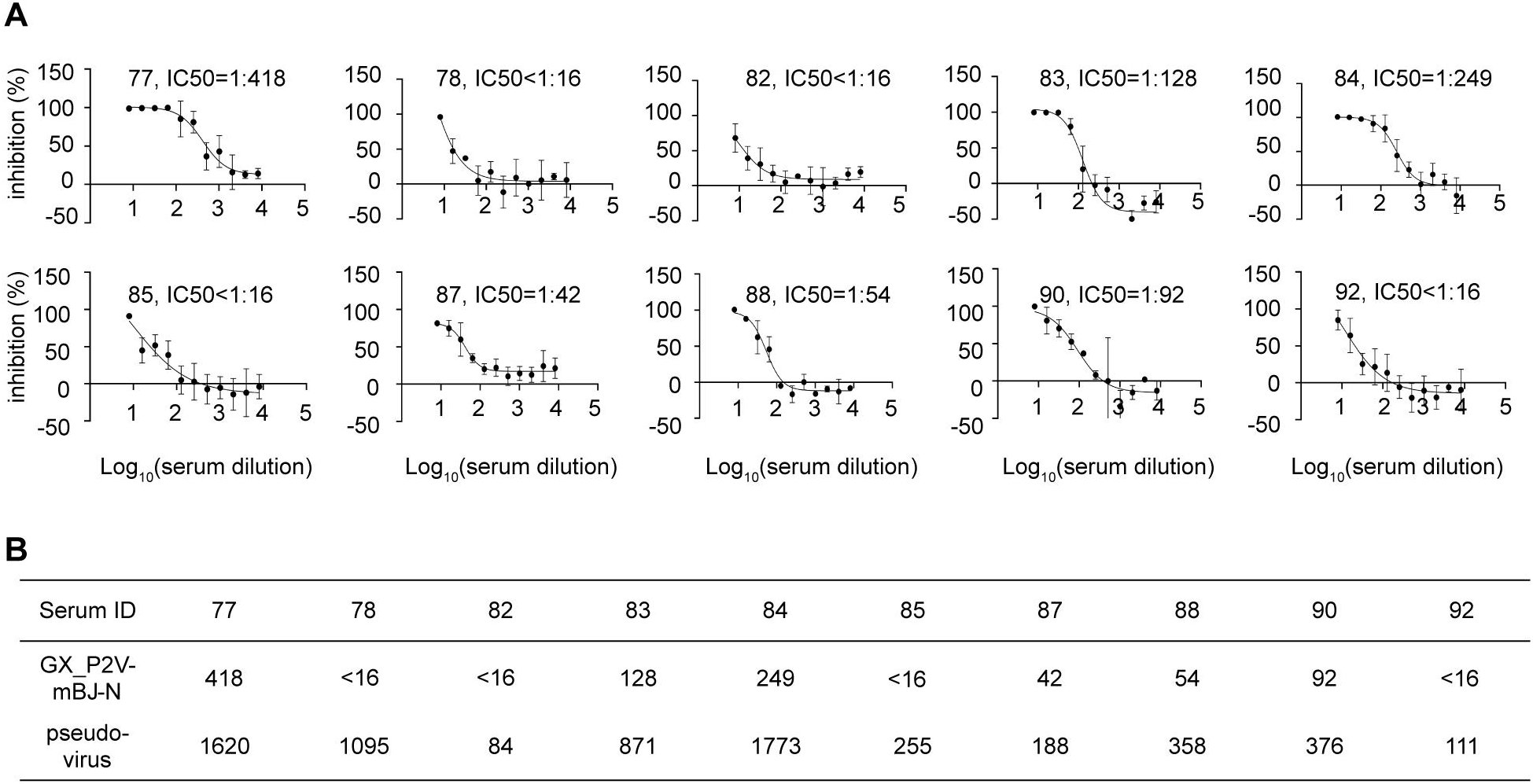
Neutralization of GX_P2V by sera from healthy blood donors. (A) Dilution-response curves for ten sera against GX_P2V-mBJ-N. Assays were performed three times; error bars show standard deviations. (B) Fifty percent neutralization titers (NT50) for the same sera measured with the VSV-based pseudovirus and with the GX_P2V-mBJ-N reporter virus.

Most sera gave the expected dilution-dependent inhibition curves against the reporter virus (Fig. 7A), and neutralizing capacity varied widely among donors: sera 77, 83, and 84 gave NT_50_ values of 1:418, 1:128, and 1:249, respectively, whereas sera 82, 85, and 92 were below the limit of detection (<1:16). More than half of the sera neutralized at dilutions above 1:40, and the GMT was 40.3. The two assays broadly agreed on rank order (Fig. 7B), but pseudovirus titers were systematically higher, with a GMT of 419, roughly 10-fold above the authentic-virus value; serum 77 gave 1,620 versus 418, serum 82 gave 84 versus 10, and serum 90 gave 376 versus 92. For an individual donor, agreement could fail altogether: Serum 78 neutralized the pseudovirus with an NT_50_ of 1,095 but did not reach 50% inhibition of the authentic virus at the lowest tested dilution (NT_50_ < 16), indicating a difference of at least 70-fold between the two assays. Ten donors are too few to describe population immunity, and the activity measured here reflects cross-reactive antibodies presumably elicited by SARS-CoV-2 infection or vaccination rather than by GX_P2V itself. The informative result here is therefore methodological: pseudovirus and authentic-virus titers are not interchangeable at the level of the individual sample, and the reporter virus provides the more direct measurement.

### Cepharanthine acts principally at the entry step

Antiviral screening likewise benefits from an authentic-virus readout, and the fluorescence of GX_P2V-mBJ-N provides a quantitative substitute for plaque reduction. We previously found CEP to be a potent inhibitor of GX_P2V in cell culture without establishing how it acts (18). To localize its target step, we combined a virion preincubation assay with time-of-addition dose-response experiments (Fig. 8). In the preincubation assay, virus was mixed with drug for 1 h at 37°C and then diluted 100-fold before infection, so that the drug concentration during infection was negligible. CEP at 25 μM had no significant effect under these conditions (Fig. 8A), which rules out direct inactivation of free virions.

**FIG 8.**
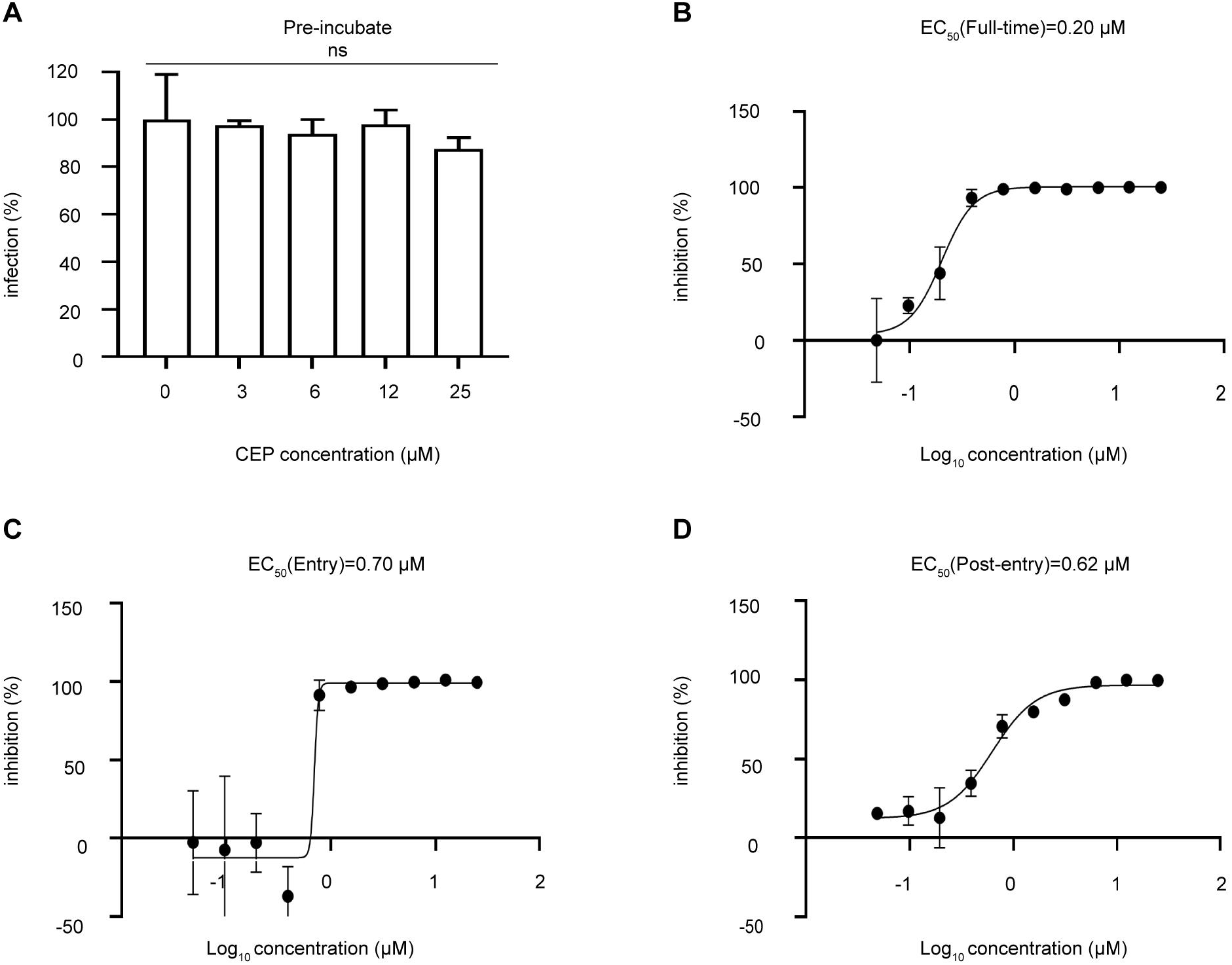
Antiviral activity of cepharanthine against GX_P2V-mBJ-N. (A) Virus was preincubated with the indicated concentrations of CEP for 1 h at 37°C, diluted 100-fold, and used to infect Vero E6 cells; fluorescence was read 48 h postinfection. Data were analyzed by one-way analysis of variance. n.s., not significant. (B to D) Concentration-response curves for CEP under full-time (B), entry-only (C), and postentry-only (D) treatment of Vero E6 cells infected with GX_P2V-mBJ-N. Fluorescence was read 48 h postinfection. Assays were performed in triplicate; error bars show standard deviations.

CEP inhibited replication in a concentration-dependent manner under all three treatment schedules (Fig. 8B to D). Inhibition was strongest when the drug was present throughout infection, with a half-maximal effective concentration (EC_50_) of 0.20 μM (Fig. 8B), compared with 0.70 μM when the drug was restricted to the entry step (Fig. 8C) and 0.62 μM when it was added only after entry (Fig. 8D). The entry-restricted curve was also markedly steeper than the other two. These values are readily explained if entry is the target. Infected cells release progeny that initiate further rounds of entry, so continuous treatment blocks the initial entry and every subsequent one, which accounts for the lower EC_50_; correspondingly, the apparent postentry activity largely reflects blockade of secondary entry, which explains why the entry-only and postentry values are nearly identical (0.70 and 0.62 μM). Taken together, these data identify entry as the principal step at which CEP acts.

## DISCUSSION

GX_P2V-mBJ-N is, to our knowledge, the first replication-competent coronavirus in which a fluorescent protein is fused to the nucleocapsid protein, and the first virus in the SARS-CoV-2-related sarbecovirus group to carry a fluorescently tagged structural protein. It is complementary to the murine hepatitis virus bearing a GFP-tagged M protein described recently (13): M labels the envelope and reports assembly and egress, whereas N labels the ribonucleoprotein core and, because N is the most abundant viral protein in infected cells, also provides a bright and quantitative readout of antigen synthesis that an M tag cannot give. The two designs converge on the same requirement, namely that the tagged structural protein must be complemented in trans by its wild-type counterpart. This suggests a general constraint on such viruses. The multifragment in vitro ligation route used here is efficient and should transfer readily to other coronaviruses.

Because N is the most abundantly expressed viral protein in infected cells, tagging it yields a correspondingly bright signal (21, 22). Earlier N-directed reporter designs avoided fusion altogether by inserting a fluorescent protein-2A cassette upstream of the N gene, so that ribosomal skipping releases the marker and leaves N intact. The genetic stability of that arrangement is unpredictable: rSARS-CoV-2/Venus-2A is stable, whereas an mNeonGreen-2A cassette at the same position in human coronavirus OC43 could not be maintained (11, 23). Direct fusion, as used here, carries a fitness cost: our virus is attenuated, no longer forms plaques, and deletes the insert within a few passages when N is not supplied in trans. In return it places the fluorophore on the virion and, in complementing cells, yields a virus that can be grown to usable titers. Interestingly, deleting the insert did not fully restore fitness: by passage 4 the insert had been reduced to a 141-bp remnant, which would be expected to give a near-wild-type phenotype, yet the virus still formed no plaques. The residual ∼47 amino acids appended to N therefore appear to impair N function, although additional adaptive mutations elsewhere in the Vero-P4 genome have not been excluded. The N terminus of N tolerates the fusion whereas the C terminus does not, which is consistent with the roles of the C-terminal domain in dimerization and in the N-M interaction during assembly. Given the readiness with which coronaviruses delete nonessential inserts, a fully stable fluorescent protein-N virus may not be achievable, and the practical alternative is the complementation scheme used here, in which intact reporter virus is produced in N-expressing cells and used within a single passage.

GX_P2V is not a human pathogen, and there is no evidence that it causes disease in pangolins (14). It is nonetheless neutralized by sera from people with no known exposure to it, which we documented previously with a pseudovirus assay in vaccinees after Omicron breakthrough infection (17) and confirm here with authentic virus. Because only ten donors were tested, these data say nothing quantitative about population immunity. What they do show is that the two assay formats are not interchangeable: pseudovirus titers exceeded authentic-virus titers about 10-fold on average and by at least 70-fold in one donor. Such divergence is expected, because a VSV pseudovirus displays only the spike protein, on a heterologous particle, and measures a single round of entry, whereas the reporter virus presents an authentic virion and reports multicycle replication; antibodies that act after entry, or that depend on native spike density and virion context, are captured only by the latter. Where neutralization data are used to infer protection, the authentic-virus measurement is the more relevant one, and GX_P2V-mBJ-N makes it practical. Separately, the attenuation of GX_P2V(short_3UTR) and the cross-reactivity documented here support continued evaluation of this strain as a live-attenuated vaccine candidate, a question that will require properly powered immunological and animal studies.

Anti-coronavirus activity of CEP has been reported by several groups. One proposal, supported by molecular docking, is that CEP binds the spike receptor-binding domain and interferes with engagement of ACE2 (24). Our preincubation result argues against a virion-directed mechanism: 25 μM CEP, more than two orders of magnitude above the EC_50_, left infectivity unchanged once the drug was diluted out. The time-of-addition data instead point to a cellular step during entry. Entry-only and postentry treatments gave similar EC_50_ values, continuous treatment gave a markedly lower one, and the entry-phase curve was steeper; all three observations follow from blockade of entry amplified across successive rounds of infection. In Vero cells, SARS-CoV-2 and related viruses enter predominantly by the endosomal route, and tetrandrine, a bisbenzylisoquinoline alkaloid closely related to CEP, blocks entry by inhibiting the two-pore calcium channel TPC2 (25). Whether CEP shares this activity is an obvious next question, for which the reporter virus is well suited.

In summary, we constructed a fluorescent, replication-competent pangolin coronavirus in which mBaoJin is fused to N, and used it to measure neutralizing antibodies against authentic virus and to assign the antiviral activity of CEP to viral entry. Two limitations should be noted. The reporter virus is genetically unstable without N complementation and must therefore be produced in N-expressing cells and used within a single passage. In addition, although a virion carrying an mBaoJin-tagged nucleocapsid is in principle suited to single-particle tracking and to live-cell imaging of assembly and egress, we did not undertake those experiments here, and the fluorescence of individual particles has not been characterized. Establishing whether GX_P2V-mBJ-N supports particle-level imaging, and combining N and M tags within one genome, are the natural extensions of this work. As it stands, the virus provides a simple authentic-virus platform for neutralization testing, antiviral screening, and mechanistic studies of sarbecoviruses under conditions that do not require high-level containment.

## MATERIALS AND METHODS

### Cell lines

Vero E6 (African green monkey kidney; ATCC CRL-1586) and BGM (buffalo green monkey kidney; CVCL_4125) cells were grown in high-glucose Dulbecco’s modified Eagle medium (DMEM; Gibco, Grand Island, NY) supplemented with 10% fetal bovine serum (FBS; HyClone, South Logan, UT) and 1% penicillin-streptomycin (Gibco) at 37°C in 5% CO_2_. N-Vero cells, a Vero E6 derivative stably expressing the SARS-CoV-2 N protein, were used for virus recovery and passage and were a gift from Qi Chen (State Key Laboratory of Pathogen and Biosecurity, Academy of Military Medical Sciences). hACE2-293T cells, transiently transfected with a SARS-CoV-2 N expression plasmid for large-scale amplification of the reporter virus, were a gift from Zheng Zhang (Shenzhen Third People’s Hospital).

### Human serum samples

Ten serum samples were obtained from healthy adult blood donors. All participants gave verbal informed consent, and the study was approved by the Ethics Committee of the 82nd Group Army Hospital of the PLA (approval no. 2025052). Samples were anonymized, were not linked to donor demographic data, and were heat inactivated at 56°C for 30 min before use.

### Cloning and assembly of full-length GX_P2V-mBJ-N cDNA

Total RNA was extracted from the supernatant of GX_P2V(short_3UTR)-infected Vero E6 cells with the FastPure viral DNA/RNA mini kit V2 (Vazyme). First-strand cDNA was synthesized with the Hifair III first-strand cDNA synthesis kit (gDNA digester plus; Yeasen) using random N6 primers and oligo(dT)_18_ at a 1:1 ratio according to the manufacturer’s protocol. Five fragments (F1 to F5) were amplified from this cDNA with PrimeSTAR GXL DNA polymerase (TaKaRa). The T7 promoter (TAATACGACTCACTATAGG) and a poly(A)_30_ tract were added by PCR at the 5′ and 3′ ends of the genome, respectively, and type IIS restriction sites (BsaI, Esp3I, and BciVI) were introduced at both ends of each fragment. Fragment boundaries were as follows: F1, T7 promoter plus nucleotides 1 to 6199; F2, nucleotides 6200 to 12397; F3a, nucleotides 12398 to 16673; F3b, nucleotides 16674 to 20953; F4, nucleotides 20954 to 24899; and F5, nucleotides 24900 to 29729 plus poly(A)_30_.

Fragments were cloned into the low-copy vector pET-40b(+) by homologous recombination with 20-bp homology arms, which limits the metabolic burden of propagating viral sequences in *Escherichia coli*; fragments 3a and 3b were cloned together in one plasmid to simplify the subsequent ligation. Plasmids were introduced into competent *E. coli* by electroporation, and transformants carrying the toxic inserts of plasmids 1 and 2 were grown at 30°C. For the reporter constructs, the mBaoJin gene was inserted upstream or downstream of the N gene and joined to it by a G4S linker, and the new plasmid was designated p5′. All subclones were verified by Sanger sequencing.

Plasmids were digested with BsaI, Esp3I, and BciVI, and the fragments were resolved on a 0.5% agarose gel, excised, and purified with the agarose gel DNA extraction kit (enhanced; TIANGEN). The full-length cDNA was assembled in two steps. Fragments were first divided into two groups (F1 to F3 and F4 to F5′) and ligated separately with T4 DNA ligase at 0.2 pmol per fragment for 24 h at 4°C. The two reactions were then combined with 2 μL of ligase, 2 μL of buffer, and 16 μL of nuclease-free water and incubated for a further 24 h at 4°C. The product was extracted twice with phenol-chloroform-isoamyl alcohol (25:24:1, pH >7.8; Meryer) and once with chloroform, precipitated with an equal volume of isopropanol at room temperature, washed twice with 70% ethanol, dried, dissolved in 10 μL of nuclease-free water, and stored at 4°C.

### In vitro transcription and virus recovery

Capped genome-length transcripts were synthesized with the mMESSAGE mMACHINE T7 transcription kit (Thermo Fisher Scientific) for 8 h at 32°C in a thermal cycler with the heated lid set to 40°C. Template DNA was removed with TURBO DNase, and the RNA was extracted with phenol-chloroform-isoamyl alcohol (25:24:1, pH <5.0; Meryer) and chloroform, precipitated with isopropanol at −20°C, washed twice with ethanol, and dissolved in 60 μL of nuclease-free water. Transcripts were electroporated immediately or stored at −80°C and used within 1 month.

N-Vero cells were seeded in a T75 flask the day before electroporation to reach about 90% confluence. Cells were trypsinized, washed in ice-cold Dulbecco’s phosphate-buffered saline (Sigma), and resuspended in 800 μL of Opti-MEM (Gibco). RNA was added, and the mixture was transferred to a prechilled 4-mm cuvette (Bio-Rad), avoiding bubbles. A single exponential-decay pulse (270 V, 950 μF) was delivered with a Gene Pulser Xcell system (Bio-Rad). Cells were seeded immediately into a T75 flask containing 10 mL of prewarmed DMEM with 10% FBS and incubated at 37°C in 5% CO_2_; the medium was replaced with DMEM containing 2% FBS 6 h later, once the cells had attached. Cytopathic effect was visible 48 h postelectroporation, and the supernatant was collected as P0. Vero E6 cells at 80% confluence were then infected with 1 mL of P0; fluorescence appeared on day 3 and increased markedly by day 4, and the supernatant was collected as P1. The virus was passaged further in N-Vero cells and in hACE2-293T cells transiently expressing N, to P11. Supernatants were aliquoted and stored at −80°C.

### RT-qPCR

Vero E6 cells were infected with GX_P2V-mBJ-N or GX_P2V-WT at an MOI of 0.01 or 5, and supernatants were collected at 0, 4, 8, 12, 24, 36, 48, and 72 h postinfection. Viral RNA was extracted according to the manufacturer’s instructions and reverse transcribed with HiScript II Q RT SuperMix for qPCR (+gDNA wiper; Vazyme). Quantitative PCR was performed with AceQ universal U+ probe master mix V2 (Vazyme), the forward primer P2V-N-F (5′-TCTTCCTGCTGCAGATTTGGAT-3′), the reverse primer P2V-N-R (5′-ATTCTGCACAAGAGTAGACTATGTATCGT-3′), and the probe P2V-N-P (5′-FAM-TGCAGACCACACAAGGCAGATGGGC-TAMRA-3′). Cycling conditions were 37°C for 2 min and 95°C for 5 min, followed by 45 cycles of 95°C for 10 s and 60°C for 30 s. Plasmid 5, which contains the P2V-N-F/P2V-N-R amplicon, was diluted 10-fold in series from 10^−4^ to 10^−11^ to generate the standard curve, corresponding to 1.17 × 10^0^ to 1.17 × 10^7^ copies per reaction.

### Western blotting

Vero E6 cells were infected with GX_P2V-mBJ-N (P1 or P4) or GX_P2V-WT at an MOI of 0.01 and lysed 48 h postinfection. Proteins were separated by SDS-PAGE and transferred to a 0.45-μm polyvinylidene difluoride membrane at a constant 100 V for 35 min in prechilled transfer buffer on ice. Membranes were blocked in 5% skim milk for 1 h at room temperature, washed three times for 5 min each in 1× TBST, incubated for 2 h at room temperature with primary antibody diluted 1:10,000, washed as before, and incubated for 1 h with secondary antibody diluted 1:5,000. After three further washes, signals were developed with SuperKine West Femto maximum-sensitivity substrate (Abbkine) and imaged on a Tanon 5200 chemiluminescence system (Tanon).

### Plaque assay

BGM cells were seeded in 6-well plates the day before infection so as to be fully confluent. Virus was diluted 10-fold in series and adsorbed by centrifugation for 1 h at room temperature. The inoculum was removed, the cells were washed twice with PBS, and each well was overlaid with 3 mL of DMEM containing 2% (vol/vol) FBS, 1% (wt/vol) methylcellulose, and 1% (vol/vol) penicillin-streptomycin. After 5 days at 37°C in 5% CO_2_, the cells were fixed in 4% paraformaldehyde for 1 h, the overlay was removed, and plaques were visualized with 0.05% crystal violet.

### Serum neutralization assay

Vero E6 cells were seeded in black 96-well plates the day before the assay to reach about 90% confluence. Sera were heat inactivated at 56°C for 30 min to inactivate complement, diluted 8-fold in DMEM containing 2% FBS, and then diluted 2-fold in series to 1:8,192. Virus was diluted to 3.2 × 10^5^ TCID_50_/mL. Equal volumes of diluted virus and serum were mixed for the test wells; serum plus medium served as the cell control, virus plus medium as the virus control, and medium alone as the negative control. Mixtures were incubated for 1 h at 37°C. Medium was removed from the plates, and 100 μL of each mixture was added per well in triplicate. After 2 h at 37°C with rocking, the inoculum was removed, the cells were washed twice with PBS, 100 μL of DMEM containing 2% FBS was added per well, and the plates were incubated for 48 h at 37°C in 5% CO_2_. Fluorescence was read on a Synergy H1 multimode plate reader (BioTek) at excitation and emission wavelengths of 479 and 520 nm. Inhibition at each dilution was calculated with the cell control wells as background (0% inhibition) and the virus control wells as 100% inhibition. Neutralization curves were generated from triplicate measurements at serial serum dilutions and fitted by nonlinear regression (four-parameter logistic model, GraphPad Prism) to obtain the 50% neutralization titer (NT_50_), defined as the reciprocal serum dilution producing 50% inhibition of fluorescence. Samples with an NT_50_ below the limit of detection (1:16) were assigned half the limit of detection (NT_50_ = 8) for statistical analyses and are indicated as <16 in figures.

### Cepharanthine antiviral assay

CEP was prepared as a 50 mM stock in dimethyl sulfoxide, diluted 2-fold in series to 1:1,024, and each dilution was then diluted 1,000-fold in DMEM containing 2% FBS to give working solutions. Four treatment schedules were used. In the preincubation group, equal volumes of 2× drug and virus were incubated for 1 h at 37°C, the mixture was diluted 100-fold and applied to cells at 100 μL per well for 2 h, and the medium was then replaced with drug-free medium. For the remaining groups, virus was diluted 100-fold to 3.2 × 10^5^ TCID_50_/mL. In the full-time group, cells were preincubated with 1× drug (100 μL per well) for 1 h, 2× drug and virus (50 μL each) were added for 2 h, and the medium was then replaced with medium containing 1× drug. The entry group was treated identically except that the medium after infection was drug free. In the postentry group, cells were infected with virus and DMEM containing 2% FBS (50 μL each) for 2 h, after which the medium was replaced with medium containing 1× drug. After 48 h at 37°C in 5% CO_2_, fluorescence was read as described above, and the EC_50_ was defined as the concentration giving 50% inhibition of viral replication.

### Statistical analysis

Statistical details for each experiment are given in the corresponding figure legends. Viral genome copy numbers are reported as means ± standard deviations of three independent experiments. Neutralization and concentration-response curves were generated from triplicate measurements and fitted by nonlinear regression to obtain NT_50_ and EC_50_ values. Differences in the preincubation assay were assessed by one-way analysis of variance; comparisons marked n.s. were not statistically significant. Analyses and plots were produced with GraphPad Prism 9.0.0.

## ACKNOWLEDGMENTS

This research was supported by the National Natural Science Foundation of China International Cooperation Research Project (NSFC-MFST, China–Mongolia) (no. 32161143027), the Beijing Natural Science Foundation Joint Fund Key Project (no. L256028), the Changping District Deputy Chief Scientist Project (no. 202506004056), the Beijing Natural Science Foundation General Program (no. 7252153), and the Key R&D Program of Xinjiang Uygur Autonomous Region (no. 2025B02022).

All the authors declare that they have no competing interests.

W.Z. and L.S. contributed to the study design; Y.M., S. Lu, and L.S. contributed to the methodology; Y.M., S. Lu, S. Luo, Y.H., C.W., X.Z., L.D., C.L., and W.C. conducted the experiments; Y.M. and S. Lu wrote the original draft; W.Z. and L.S. reviewed and edited the manuscript; W.Z. and L.S. acquired funding; S. Luo, Y.H., C.W., X.Z., L.D., C.L., and W.C. provided resources; W.Z. and L.S. supervised the study. All authors have read and approved the final manuscript.

